# Improved Metabolic Flux Estimations through Compositional Data Analysis

**DOI:** 10.64898/2026.08.07.742769

**Authors:** Alberte Sloth Carlsen, Te Chen, Nicholas Luke Cowie, Christian Brinch, Teddy Groves, Lars Keld Nielsen

**Author notes:** All authors contribute equally (Alberte Sloth Carlsen, Te Chen).

## Abstract

Isotopic Metabolic Flux Analysis (I-MFA) is a standard approach for estimating intracellular metabolic fluxes. I-MFA infers fluxes by comparing simulated and measured metabolite isotopologue distributions (MIDs) of metabolites from isotope labeling experiments. MIDs represent fractional abundances that strictly sum to one for any given metabolite, thus they are inherently compositional data. However, state-of-the-art estimation approaches rely on calculating standard Euclidean distances between MIDs in a non-compositional paradigm, introducing a systemic bias. To resolve this, our study proposes compositional I-MFA. We demonstrate how to construct a meaningful orthonormal basis for MIDs via ordered sequential binary partitioning, which can be used to perform isometric log-ratio (ILR) transformation. As a minimal change to existing I-MFA workflows, we suggest estimating fluxes by minimizing Euclidean distances between ILR-transformed MIDs. We validated this framework against traditional methods using both a toy model and a biologically realistic model, evaluating point estimates, sensitivity across varied true fluxes, and confidence intervals. In the two examples, compositional I-MFA consistently outperformed traditional approaches, reducing mean squared error of flux point estimates by an average of 42.6% and substantially narrowing confidence intervals. We conclude that compositional data analysis significantly improves I-MFA and can be implemented as a simple drop-in replacement for current pipelines.

**Graphical Abstract:** 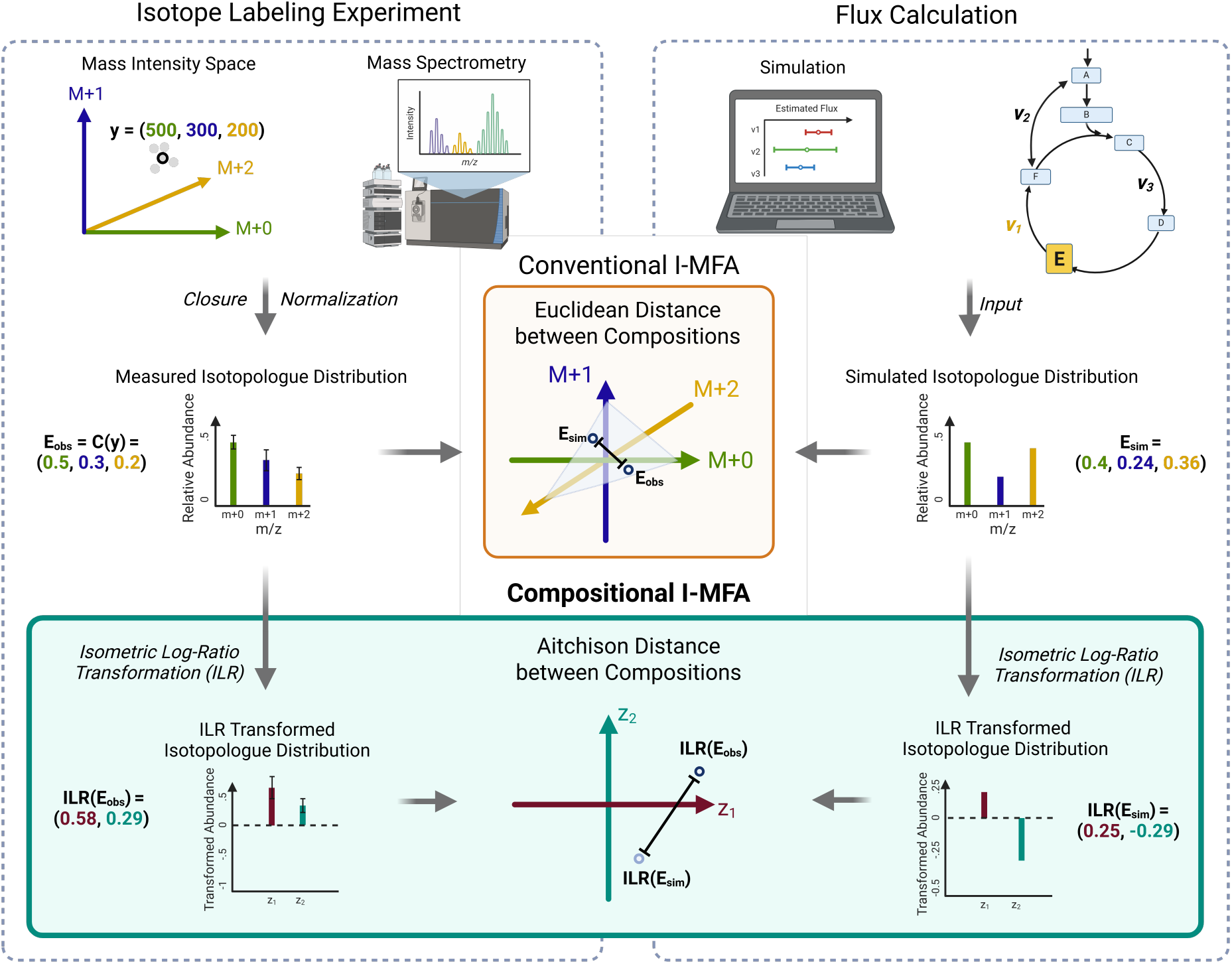

**Highlights:**

- New compositional data approach improves metabolic flux estimation.
- This data transformation requires minimal changes to existing workflows.
- The new method reduced MSE of flux estimates by 42.6% in two examples tested.
- The confidence intervals of the estimated fluxes were substantially narrowed.
- Estimation accuracy remained robust across a wide range of metabolic fluxes.

## 1. Introduction

The metabolic phenotype of a cell is defined by the distribution of fluxes through its metabolic network. Unfortunately, these fluxes can generally not be measured directly or inferred from exchange rates with the environment.

Isotopic metabolic flux analysis (I-MFA) solves this problem by exploiting isotope labeling experiments, where cells are cultivated in the presence of known quantities of positionally isotope-labeled substrates (Wiechert and Nöh, 2021). Cell metabolism predictably redistributes these labeled molecules, so that it is possible to simulate the measurable isotopic distribution of a cell’s metabolites given its flux state. Given a simulator and a set of measurements, I-MFA infers a flux state with a simulated isotopic distribution that accurately approximates the measured isotopic distribution.

Isotopic distributions can be measured using nuclear magnetic resonance (NMR) spectroscopy or mass spectrometry (MS). While NMR provides highly detailed positional information (isotopomers), MS is more commonly used due to its higher sensitivity (Gowda and Raftery, 2021). Measuring intact ions, standard MS cannot distinguish between positional isotopomers, as they share identical atomic masses. This work focuses on the latter case, where the primary unit of measurement is a metabolite isotopologue distribution (MID, also referred to as “mass isotopomer distribution”), i.e. a non-negative unit vector specifying the relative abundance of a compound’s isotopologues (Bluck, 2013). Note that the arguments made here for isotopologues are equally valid for finer-grained NMR isotopomer measurements.

MIDs are preferred over absolute measurements because they represent relative abundances, which inherently normalize for variations in sample concentration, extraction efficiency, and instrument sensitivity. MIDs are also used during natural isotope abundance correction and simulations. Most I-MFA studies also supply MIDs as their data for scientific reporting. Because MIDs are non-negative and sum to one, they are examples of compositional data. The statistical treatment of compositional data requires a specialized framework known as Compositional Data Analysis (CoDa). Although pioneered in the geosciences for evaluating geochemical and meteoritic compositions (Aitchison, 1982; Pawlowsky-Glahn and Egozcue, 2006), CoDa has since been widely adopted in other scientific domains. In metagenomics, for example, researchers increasingly rely on CoDa to analyze shifts in microbial species compositions while avoiding the spurious correlations caused by standard statistical methods (Gloor et al., 2017; Quinn et al., 2018).

To quantify the discrepancy between a measured and simulated MID, most I-MFA analyzes have used the component-wise weighted sum of squared residuals (SSR) directly in Euclidean space without considering compositional constraints. While previous studies (Kaste and Shachar-Hill, 2024) have noted the inconsistencies of SSR in I-MFA, no formal mathematical solution has been proposed. Several software packages implement I-MFA workflows (Rahim et al., 2022; Stratmann et al., 2025; Sokol et al., 2012; Quek et al., 2009) and all rely on Euclidean metrics (e.g., SSR) to compare MIDs. To date, no tool or study utilizes a metric that satisfies the necessary compositional properties.

Here, we demonstrate that the traditional approach creates a geometric mismatch: while MIDs are constrained to a simplex, SSR assumes an unconstrained space. To address this, we advocate for the use of CoDa, especially log-ratio transformations that map the simplex onto unconstrained real space. Using examples, we demonstrate that comparing MIDs using non-compositional methods leads to poorer I-MFA performance compared with compositional methods. Furthermore, we show that it is straightforward to extend standard I-MFA workflows to include these methods.

## 2. Materials and methods

Our main hypothesis is that I-MFA can be improved by using compositional data analysis to compare measured and simulated isotopologue distributions. See section Appendix A and Appendix B for background knowledge. To test this, we implemented a compositional I-MFA workflow and compared it with traditional I-MFA on two well-studied test problems.

### 2.1. Proposed compositional I-MFA method

Our compositional I-MFA workflow aims to diverge as little as possible from traditional I-MFA without applying non-compositional data analysis methods to compositional data. Our workflow therefore differs from traditional I-MFA only in how discrepancies between measured and simulated MIDs are quantified. Whereas traditional I-MFA applies Euclidean objective functions to MIDs directly, we propose to do so after log-ratio transformations. As a concrete example of this strategy, we propose applying the isometric log ratio transformation (ILR) to both measured and simulated MIDs and then minimizing the SSR on the transformed values. To help with interpretation, we also propose a procedure for constructing sequential binary partitions that produces interpretable results when applied to MIDs.

#### 2.1.1. ILR transformation for MIDs

Considering that MIDs are compositional, it is important to preserve the compositional properties explained in Appendix B.5. This can be achieved using a log ratio transformation that maps a composition from its native simplex to an unconstrained Euclidean space, see Supplementary Fig. B.6. The Isometric Log Ratio (ILR) transformation is a popular choice, but requires an appropriate orthonormal basis of the target simplex (Pawlowsky-Glahn and Egozcue, 2006). For an explanation of ILR and the related centered log ratio transformation, see Appendix B.2.

To construct an orthonormal basis for the *D*-part MID **x** = *x*_*m*+0_, *x*_*m*+1_, …, *x*_*m*+(*D*−1)_ with *D* mass shifts, we propose an ordered sequential binary partitioning method.

The first row of the basis separates the lightest isotopologue *x*_*m*+0_, indicated by the number 1, from all heavier isotopologues, indicated by the number −1. The second row assigns *x*_*m*+0_ the number 0, then separates *x*_*m*+1_ from the heavier isotopologues in the same way as in the first row. Subsequent rows repeat this pattern until the partition is complete in row *D* − 1.

For example, for a four part MID the ordered sequential binary partition is as follows:

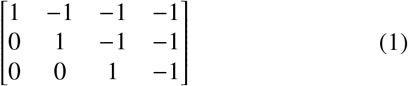

Applying the normalization method in Supplementary Equation B.6 to obtain an orthonormal basis **Ψ**, the *i*-th component of the ILR-transformed version of **x** is then as described in Dumuid et al. (2017):

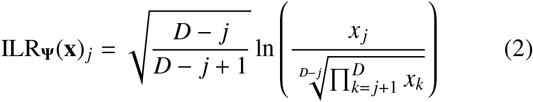

Using our method, each value of ILR_**Ψ**_(**x**)_*i*_ has a natural interpretation, representing the relative abundance of the *j*-th isotopologue compared with heavier isotopologues.

#### 2.1.2. Proposed objective function

Instead of minimizing the SSR between the raw simulated and measured isotopologue distributions, we propose to minimize the SSR between the ILR-transformed simulated MID and the ILR-transformed measured MID. We define this compositional objective function as SSR_ILR_:

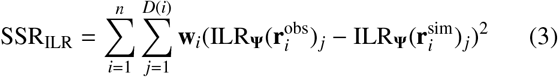

where **Ψ** is an orthonormal basis, *n* is the number of metabolites, 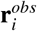 is the observed composition with length *D*(*i*) of the *i*^*th*^ metabolite. 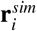 is the simulated composition. **w**_*i*_ is a vector of non-negative weights with length *D*(*i*), calculated from the measurement error vector *σ*_*i*_ of the ILR transformed measured MID:

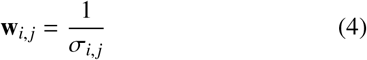

Note that our objective function has the same relationship with the Aitchison distance as defined as in Supplementary Equation B.8 as SSR has with the Euclidean distance (see Supplementary Equation B.7).

### 2.2. Comparison Experiments

To evaluate the benefits of compositional I-MFA, we copied two commonly used, previously published, example reaction networks. We conducted three different simulation experiments using compositional I-MFA to compare our results with those of traditional I-MFA methods.

The first toy reaction network example (toy network) comes from Antoniewicz et al. (2006) and Antoniewicz et al. (2007). Both papers used the same example reaction network to showcase the method for calculating flux confidence intervals and the performance of elementary metabolite unit (EMU) method. Using this network, we could directly compare our results with the original I-MFA methods. The second TCA cycle reaction network (TCA cycle) comes from Quek et al. (2009) and is a more realistic biological example using data from a real I-MFA experiment.

We performed three experiments on each of these two models. In the first experiment, we used original data from Antoniewicz et al. (2007) and Quek et al. (2009) to calculate ground truth flux, then simulated measurement errors and fit both compositional and traditional I-MFA to the perturbed data. This was done to ensure that the proposed compositional I-MFA works in a basic setting. Our second experiment compared three types of flux confidence intervals calculated using compositional I-MFA with the same confidence intervals calculated using traditional I-MFA. Lastly, to test whether compositional I-MFA maintains accurate flux estimation across a range of realistic flux values, we carried out a sensitivity analysis by systematically varying ground truth fluxes.

#### 2.2.1. Example reaction networks

We evaluated our approach using two previously published reaction networks, schematized in Figs. 1 and 2.

**Fig. 1:**
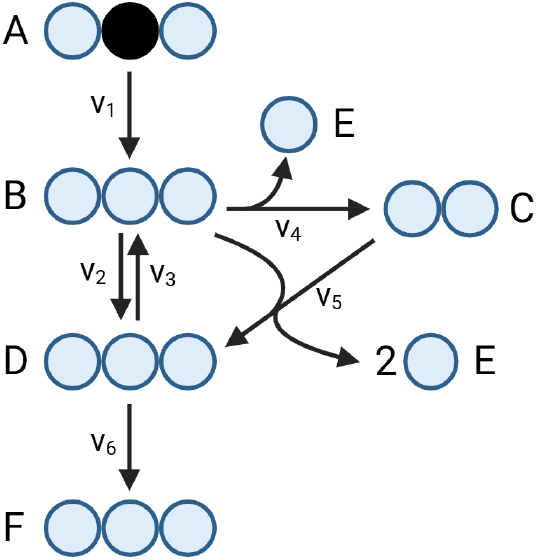
Toy example reaction network. The network has 6 metabolites and 6 reactions. Each metabolite has maximum 3 labelable atoms. Metabolite A is fully labeled at position 2 and *v*1 is fixed at 100 in all experiments. **Antoniewicz et al. (2007)**

**Fig. 2:**
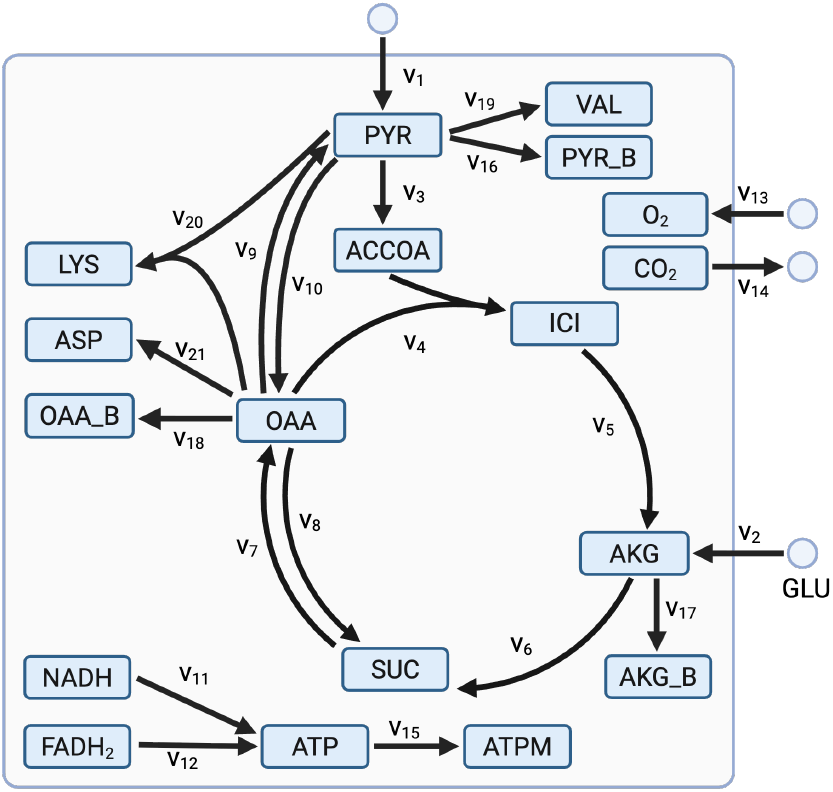
TCA cycle reaction network. This network represents the tricarboxylic acid cycle in central metabolism. The network has 21 reactions and 15 metabolites, including 7 sink reactions and 4 boundary reactions. **Quek et al. (2009)**

The first network consists of six metabolites, four irreversible reactions, and one reversible reaction (represented as *v*2 and *v*3). In all experiments, we assumed that metabolite A is fully labeled at position 2 and the external flux *v*1 is fixed at 100. The aim of this analysis is to infer the unknown fluxes *v*2, …, *v*6 using a single MID measurement of metabolite F. In the ground truth experiment and confidence interval experiment, we fixed fluxes **v** = [100 110 50 20 20 80] as the solution, so that the true MID for metabolite F is the four-part composition [0.0001 0.8008 0.1983 0.0009]. We set *v*3 and *v*5 in this network as independent fluxes. To produce comparable results, these values come from Section 3.1 in Antoniewicz et al. (2007) and Section 2.3 in Antoniewicz et al. (2006). The TCA cycle in the second example (Fig. 2) comes from Quek et al. (2009), which simplifies the tricarboxylic acid cycle in central metabolism. The network has 21 reactions and 15 metabolites, including 7 sink reactions and 4 boundary reactions. In this experiment, the label input mixture was 50% fully labeled pyruvate ([U-^13^C] Pyr), 50% position 1 labeled pyruvate ([1-^13^C] Pyr) and 100% position 1 labeled glutamate ([1-^13^C] Glu). The pyruvate boundary flux *v*1 was set to 1 in all experiments. The simulated measurements are the MIDs for the metabolites valine (Val), lysine (Lys), aspartate (Asp) and succinate (Suc). The true value of these can be seen in the Supplementary Table D.6.

Full reaction lists, atom transitions, and specific network parameters for both models are provided in Appendix D.

#### 2.2.2. Independent fluxes and free fluxes

To constrain the mass balance of metabolites in the networks, we define the stoichiometric matrix **S** ∈ ℝ^*M*×*N*^, where *M* is the number of internal metabolites and *N* is the number of reactions. Under the metabolic steady-state assumption, the net accumulation of all intracellular metabolites is zero:

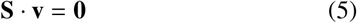

where **v** is the vector of fluxes and **0** is a null vector.

Because metabolic networks are typically underdetermined (*M* < *N*), the system possesses fewer mass balance constraints than unknown reactions. The solution space is therefore parameterized as a linear combination of a reduced set of independent fluxes, **v**_ind_:

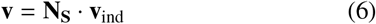

where **N**_**S**_ is a basis matrix for the null space of **S**. The dimension of **v**_ind_ is equal to the nullity of **S**.

In our network models, a subset of these independent fluxes is constrained by fixing the known input and output fluxes (e.g., substrate uptake reaction and biomass reaction). The remaining unconstrained variables in **v**_ind_ constitute the *free fluxes*, which are the variables explicitly estimated during the optimization process. For the Toy network, the free fluxes are *v*_3_ and *v*_5_; for the TCA cycle network, they are *v*_2_, *v*_8_, and *v*_9_.

#### 2.2.3. Simulated MID measurement sets

In standard I-MFA workflows, MIDs are derived from mass spectrometry (MS) data, where the raw signals are recorded as ion abundances (e.g., integrated peak areas) for each isotopologue. These raw abundances are subject to multiplicative instrumental noise before being normalized into fractional abundances. To rigorously evaluate the objective function landscape, it is necessary to model the data generating process at the level of these pre-normalized abundances.

For each network, we generated 10000 random abundance measurements based on the ground truth MID values using a log-normal model:

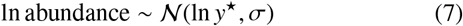

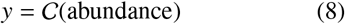

where *y* is the observed MID, *y*^⋆^ is the ground truth MID, *σ* is a vector of log-scale measurement errors, *C* represents the closure operation defined in Supplementary Equation B.2 and *N* represents the normal distribution. While raw MS abundances scale with the total metabolite pool size, the scale invariance of the closure operator allows us to apply the log-normal noise directly to the fractional *y*^⋆^ without loss of generality.

Our choice of a log-normal distribution is motivated by classical models of analytical measurement error, which demonstrate that MS ion abundances exhibit an approximately constant coefficient of variation (CV) across the<u>ir li</u>near dynamic range (David M. Rocke, 2026). We set 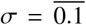, corresponding to approximately 10% coefficient of variation on the raw abundances, as this generated maximum standard deviations of approximately 2% when pushed forward onto MIDs, which is consistent with a realistic approximate best case scenario for quantitative mass spectrometry: see (Heuillet et al., 2018; Antoniewicz, 2018) for discussion of realistic errors on isotopologue distributions under MS analysis.

This measurement model is a simplification: real MS measurements are subject to additional sources of error and bias such as saturation of the ion detector at high intensities, reduced accuracy for low-intensity transitions as the target peak approaches the scale of background oscillations and ion suppression (Lu et al., 2017; Evard et al., 2016; Zhou et al., 2017). However, these instrument-specific errors are independent of the mathematical choice between compositional and non-compositional I-MFA frameworks, and are thus outside the scope of this study.

#### 2.2.4. Ground truth flux estimation

We fit each simulated measurement set using our own implementations of traditional and compositional I-MFA. For traditional I-MFA, we estimated fluxes by minimizing the SSR between the simulated and observed MIDs. For compositional I-MFA, the objective function minimized the SSR of the ILR-transformed simulated and observed distributions using Equation 3. In both networks, the reversible fluxes are divided into forward and reverse. Therefore, all fluxes are strictly positive. To ensure this, we optimized the log-transformed fluxes and back-transformed before calculating the simulated MIDs.

To evaluate estimation accuracy across the 10000 groundtruth simulations, we calculated the squared error between each estimated free flux and its known ground-truth value. To account for scale differences between distinct reactions, squared errors were not aggregated across the network; all averages are reported with respect to specific parameterizations and simulations.

#### 2.2.5. Confidence interval estimation

To estimate parameter confidence intervals (CIs) for both methods, we employed Monte Carlo simulations as described by Antoniewicz et al. (2006). The Monte Carlo 95 % CI was defined by the 2.5% and 97.5% quantiles from the 10000 estimations. We compared these against two alternative estimation methods: profile likelihood and Fisher Information Matrix (FIM). Profile likelihood CIs were calculated by fixing the parameter of interest at sequential values, re-optimizing the remaining free parameters, and observing the likelihood profile using `pyPESTO`(Schälte et al., 2026). For the FIM-based approach, we obtained a linear approximation Σ of the flux covariance matrix by projecting the inverse Hessian of the free fluxes:

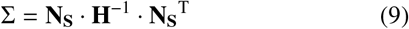

where **H** is the Hessian matrix, indicating the local curvature of the objective function under small changes in measurement values and **N**_**S**_ is the null space of the stoichiometric matrix. The CI for flux *i* is then calculated as

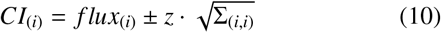

where Σ_(*i,i*)_ is the variance of flux *i* obtained from the covariance matrix, and *z* is the critical value corresponding to the desired confidence level. The profile likelihood and the FIM are calculated with random drawn simulated measurements and the variance from the simulated measurements of MIDs.

### 2.3. Sensitivity analysis

To evaluate method robustness across the parameter space, we supplemented the 10000 ground-truth datasets with an evenly spaced grid of free fluxes ranging from 50% to 150% of the true flux values (see Appendix D for network-specific boundaries). For each cell in this flux grid, we generated 50 random MS measurement sets using the log-normal model described above, allowing us to map the convergence properties of the estimators.

#### 2.3.1. Implementation and optimization algorithms

For efficient simulation of isotopologue distributions given fluxes, we re-implemented the EMU algorithm from Antoniewicz et al. (2007). For gradient based minimization we used the BFGS optimiser provided by the Python library optimistix (Rader et al., 2024). All code used to generate our results, as well as reproduction instructions, is available at https://github.com/dtu-qmcm/cmfa.

## 3. Results

### 3.1. Better ground truth flux estimation

Based on the 10000 simulated MIDs, we calculated the residuals between the true and estimated fluxes. The residuals for the free fluxes in the Toy network and the TCA cycle are shown in Fig. 3, along with their 95% predictive intervals. In our experiments, the mean squared errors (MSE) of the residuals for the free fluxes were consistently lower for compositional I-MFA compared to traditional I-MFA. These MSE values are also presented in Fig. 3. For the Toy network, the MSE for *v*_3_ was reduced from 181.13 to 142.17 using compositional I-MFA, which is a 21.5% reduction. For the TCA cycle example, the most significant improvement occurred for *v*_9_, where the MSE decreased by 70.1% compared with traditional I-MFA. For a summary of all fluxes see Supplementary Tables E.7-E.10.

**Fig. 3:**
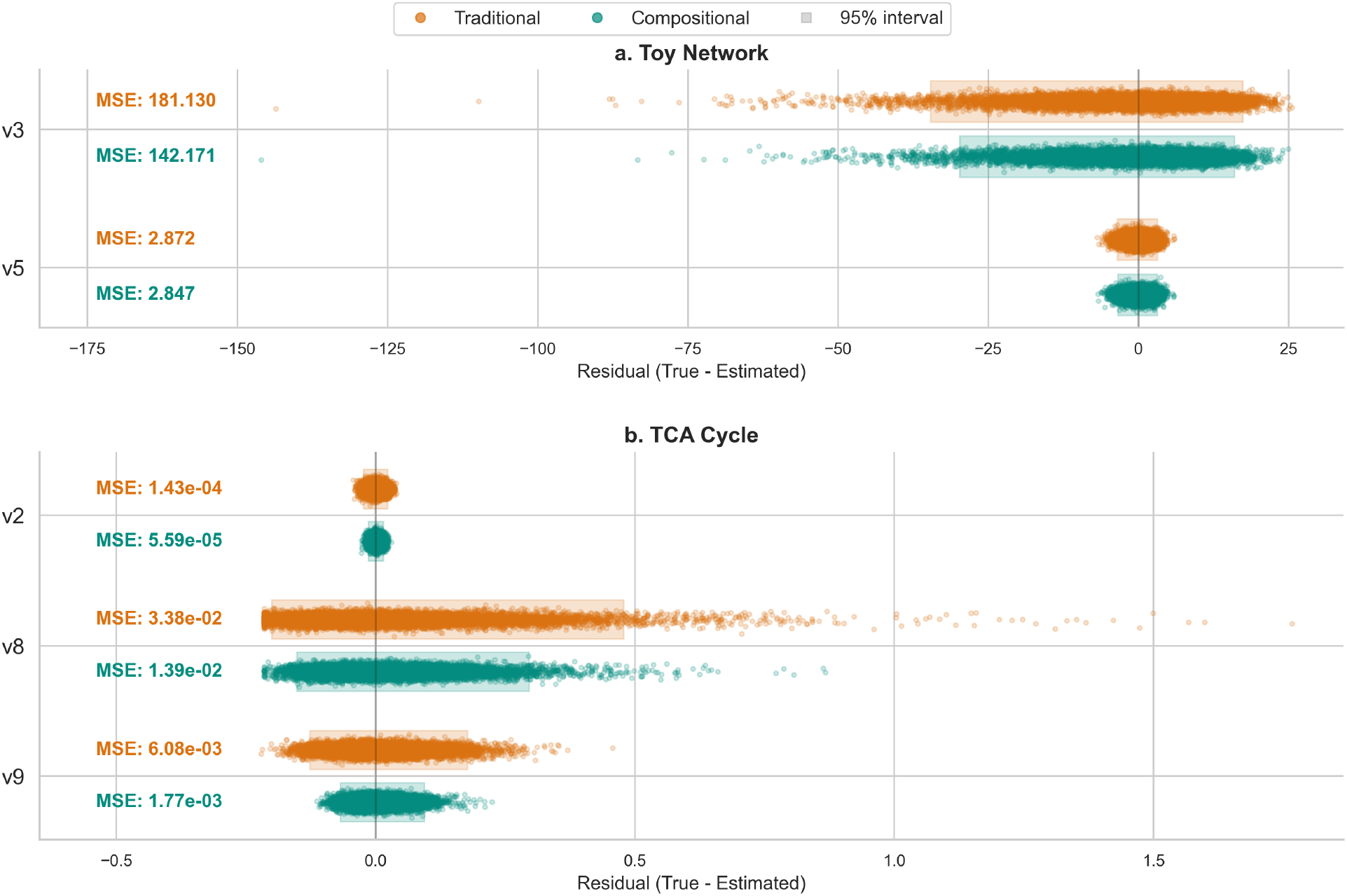
Free flux residuals and predictive performance. Comparison of residuals (True Flux-Estimated Flux) for metabolic fluxes in the **a**. Toy network and **b**. TCA cycle. Each distribution consists of *n* = 10000 data points, representing individual estimation errors for traditional I-MFA (orange) and compositional I-MFA (cyan). Shaded boxes indicate the 95% predictive intervals, and Mean Squared Error (MSE) values for each method are annotated on the left. The vertical gray line at zero represents perfect estimation. Compositional I-MFA consistently achieves lower MSE and narrower residual distributions across most fluxes

### 3.2. Tighter confidence intervals on ground truth flux

Compositional I-MFA improved the precision of the estimated confidence intervals across all three methods used in Antoniewicz et al. (2006). The 95% CIs derived from Monte Carlo simulation for the free fluxes were the tightest of the three methods, particularly in the TCA cycle example. The distributions of these Monte Carlo 95% CIs are illustrated as histograms in the top row of Fig. 4. For comparison, the probability densities calculated using FIM are shown as curves in the middle row of Fig. 4. The bottom row of Fig. 4 shows the profile likelihoods, where the intersection of the curves with the threshold defines the upper and lower bounds of the 95% CI.

**Fig. 4:**
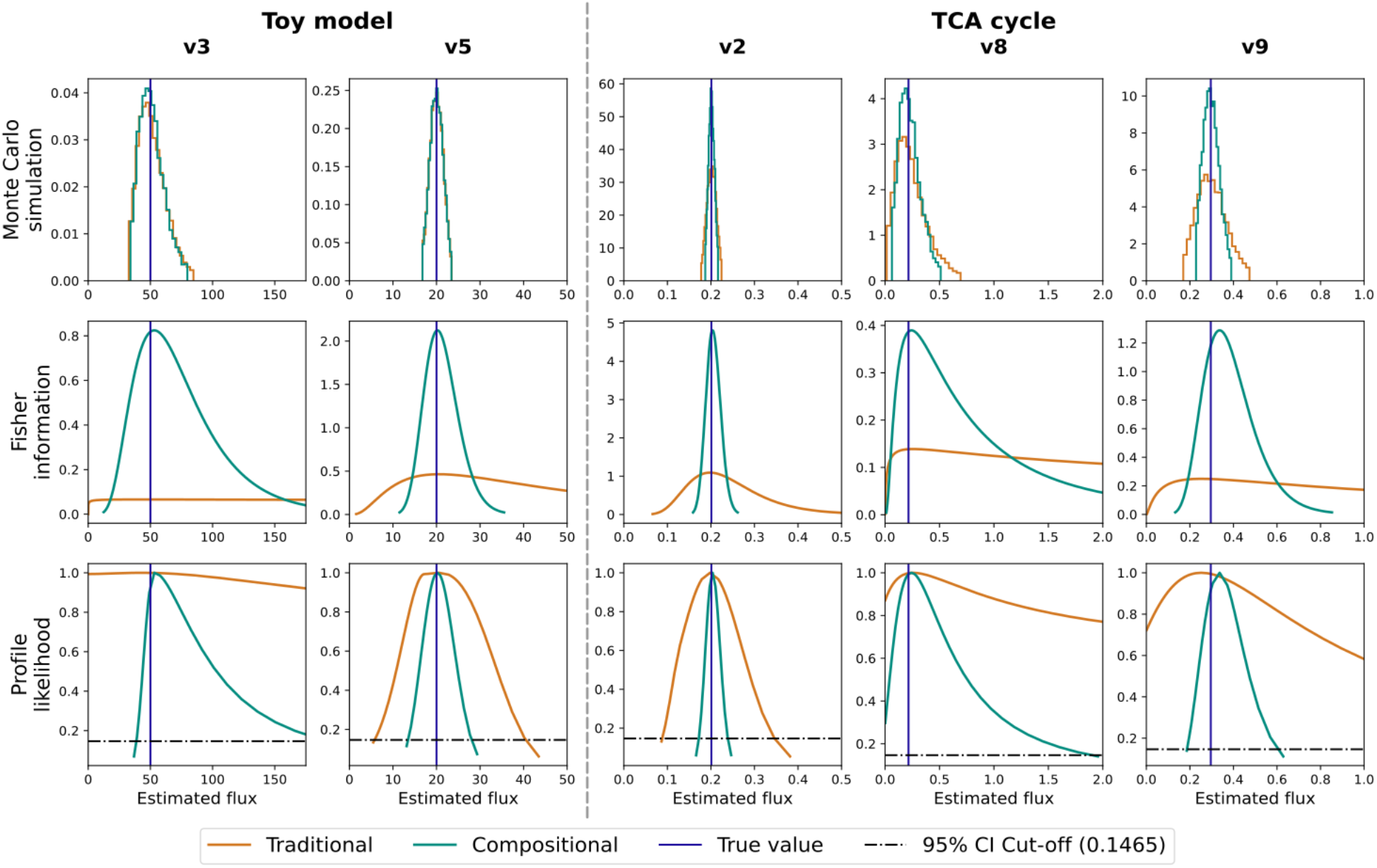
Comparison of uncertainty quantification. From top to bottom the rows represent the Monte Carlo simulation, FIM and profile likelihood. In each figure orange represents traditional I-MFA and cyan represents compositional I-MFA. The blue vertical lines mark the true flux values, and the black horizontal lines represent the profile likelihood cutoff threshold. The histograms in the first row show the estimates from 2.5% quantile to 97.5% quantile through Monte Carlo simulations. The curves in the middle row are the probability densities based on FIM. The bottom row shows the profile likelihoods. The profile likelihood 95% CIs are the points where the lines meet the horizontal cutoff in black dashed line. The figures x-axis range are truncated and the full 95% CIs can be seen in Supplementary Table E.11.

Across all evaluated free fluxes, the traditional framework consistently shows wider or structurally unbounded profiles, as shown in Fig. 4 and Supplementary Table E.11. For instance, in the Toy network, the traditional I-MFA FIM-based 95% CI for flux *v*3 spanned an unrealistically wide range of [0.0003, 5 921 400.0000]. Under the compositional I-MFA framework, this interval was narrowed to a realistic bound range of [20.5030, 137.6500]. Similarly, for flux *v*9 in the TCA cycle example, the FIM-based 95% CI shrank from [0.0109, 5.8193] to [0.1835, 0.6177]. CIs calculated based on profile likelihoods show the same trend where compositional I-MFA showed consistently narrower intervals compared to the traditional method, see Fig. 4 and Supplementary Table E.11. Additionally, the traditional methods profile likelihood CIs of some fluxes, such as *v*3 in Toy Network, and *v*8 in TCA cycle, did not reach the threshold within the predefined lower and upper bounds of 0.0009 and 1096.6000 respectively, while compositional I-MFA yields most intervals in bounds.

### 3.3. Sensitivity analysis shows robustness across wide range

The example of ground truth prediction showed higher accuracy with various simulated MIDs, however it did not test how robust compositional I-MFA is. A sensitivity analysis with an evenly spaced grid of flux sets was used to explore its robustness. For each flux set, the difference between the MSE of the traditional and compositional methods was calculated. Positive values indicate that compositional I-MFA is more accurate than traditional I-MFA. In the Toy network example (Fig. 5a), the MSE for compositional I-MFA was lower in the majority of cases, with 114 and 70 out of 121 for *v*_3_ and *v*_5_, respectively. The mean of the difference is 59.26 for *v*_3_ and 0.03 for *v*_5_. In the TCA cycle example, compositional I-MFA yielded a lower MSE in all instances as shown in Fig. 5b. In this figure, the points are ordered by their corresponding flux values; thus points on the same line share the same true flux values, ranging from −50% to +50% compared with the ground truth flux.

**Fig. 5:**
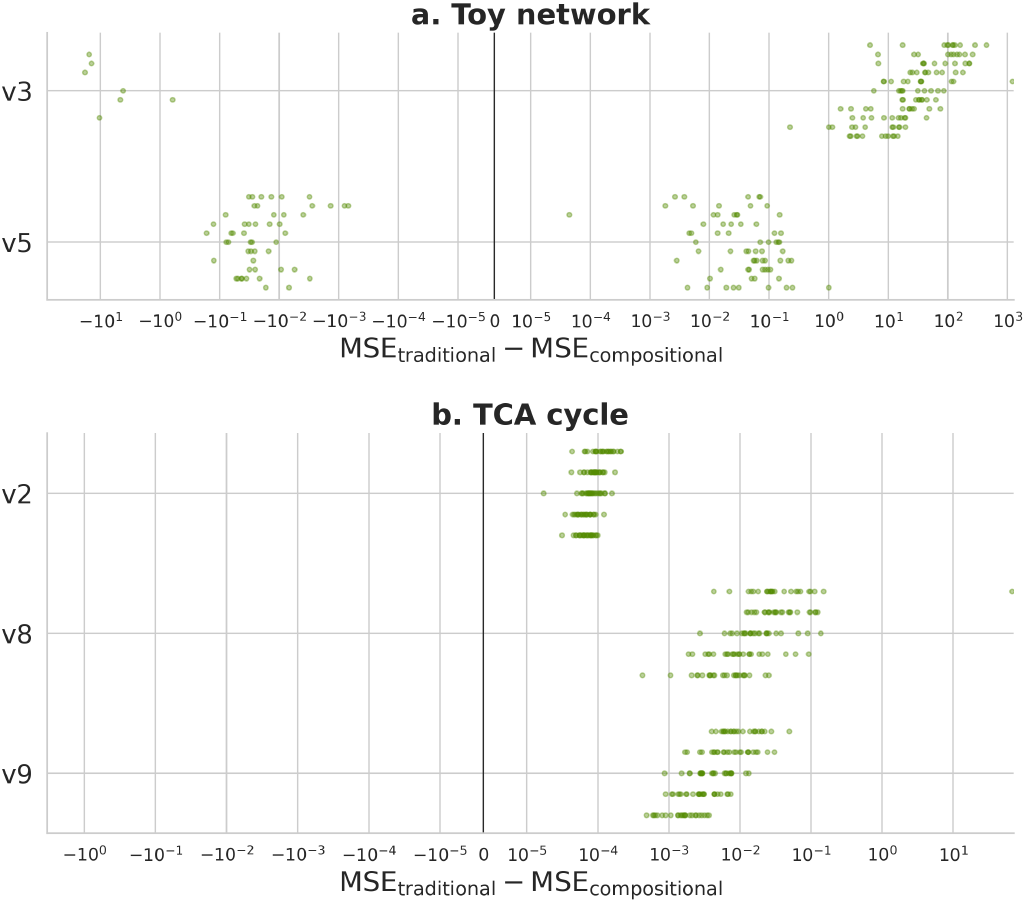
Difference in MSE between traditional and compositional I-MFA. (*MS E*_*traditional*_ − *MS E*_*compositional*_) across varying fluxes for **a**. the Toy network and **b**. the TCA cycle. Positive values denote lower error for the compositional method. Data points are distributed along the y-axis corresponding to their true flux values (Toy network: *v*3 ∈ [25, 75], *v*5 ∈ [10, 30]; TCA cycle: *v*2 ∈ [0.1005, 0.2015], *v*8 ∈ [0.1075, 0.3235], *v*9 ∈ [0.148, 0.444]). The mean of the differences is *v*_3_: 59.2630, *v*_5_: 0.0304, *v*_2_: 0.0001, *v*_8_: 0.5668, *v*_9_: 0.0065. The x-axis uses a symmetric log scale with a linear range from − 10^−5^ to 10^−5^.

## 4. Discussion

### 4.1. Compositional I-MFA yields more accurate flux estimations

By fundamentally correcting how MIDs are handled in I-MFA, compositional data analysis resolves a long-standing systemic bias in I-MFA. Compositional I-MFA consistently yields more accurate flux estimates than traditional I-MFA in both of our examples, reducing MSE of flux estimates by an average of 42.6% and substantially narrowing confidence intervals. Importantly, systematically varying the ground truth demonstrates that the improvement from using CoDa is robust across a wide range of metabolic fluxes. Because our proposed method is computationally simple and requires minimal modifications to existing software pipelines, it serves as a seamless, drop-in replacement for any workflow that requires comparing simulated and observed MIDs.

### 4.2. Estimation of confidence intervals

Quantifying the uncertainty around estimated fluxes presents well-documented challenges. Antoniewicz et al. (2006) demonstrated that local linearization using the FIM yields unreliable intervals in traditional I-MFA. While they proposed an alternative profile-likelihood-like method that drastically reduced computational time relative to brute-force grid searches, their approach still requires an iterative algorithm executed independently for each flux parameter.

In this work, we evaluated local linearization via the FIM across both the traditional and compositional I-MFA frameworks, and compared them against profile likelihoods and

Monte Carlo simulations. Our experiments corroborate the conclusion of Antoniewicz et al. (2006) that FIM produces unfeasible confidence intervals in traditional I-MFA. However, applying the FIM in our compositional I-MFA framework yielded tightly bounded confidence intervals. Thus, under compositional I-MFA, FIM restores its scientific utility due to its conceptual simplicity, fast computational time and accuracy.

### 4.3. Limitations of our analysis

While we recommend our approach for comparing simulated and observed MIDs, it is important to note that it is possible, and in some ways preferable, to perform I-MFA without such comparisons. Alternatively, one can construct a full forward simulation model where a flux set is pushed forward to a mass spectrometer output, i.e. a number of reads or area under the curve. In this alternative framework, observed MIDs are by-passed entirely, while simulated MIDs appear merely as an intermediate computational stage between the flux set and the expected MS readouts. As well as not requiring compositional data analysis, since MS outputs are not compositions, this approach also has the benefit of being able to model explicitly MS-specific phenomena such as upper and lower detection limits and dilution-related heteroskedasticity. On the other hand, a full forward simulation model is only feasible if unclosed MS outputs are available, which might not be the case depending on the analytics workflow or the approach to natural isotope abundance correction. We therefore consider that our proposed approach and full forward simulation are both valid options for modeling mass spectrometry measurements in the context of I-MFA.

In this study, we evaluated our framework using simulated measurement data with fully resolved, non-zero values. However, real-world experimental MIDs frequently contain zeros, which present a unique challenge in CoDa due to the log-ratio operation. These zeros typically arise either as essential zeros (where an isotopologue is biologically absent) or due to detection limits of the mass spectrometer. Standard CoDa strategies to address this include replacing zero values with small imputed values or omitting components containing zeros entirely (Pawlowsky-Glahn and Egozcue, 2006). While both strategies are readily applicable to the I-MFA workflow, evaluating their specific impacts on flux estimation error is beyond the scope of this initial proof-of-concept study. Because zero-handling introduces an independent layer of measurement artifact manipulation, we deliberately isolated our evaluation to clean datasets to clearly establish the core performance of the compositional paradigm.

For simplicity we did not consider Bayesian I-MFA (Theorell et al., 2017), where the comparison of observed and simulated MIDs is embedded in a Bayesian statistical model that specifies the probability density *p*(*r*^obs^, **v**, *θ*) for any possible observation *r*^obs^, flux **v** and arbitrary parameter vector *θ*. We expect that compositional I-MFA will be especially beneficial in the context of Bayesian statistical modeling, as it ensures that positive probability mass is assigned only to physically possible data configurations while avoiding a discontinuous likelihood function. Thus, key components of Bayesian workflow like posterior and prior predictive checking and simulation based calibration (Gelman et al., 2026) are available and the posterior distribution can be explored by gradient-based MCMC samplers that require local smoothness.

This study aims to present the compositional data analysis framework rather than implement the method in software. We hope to inspire the community to expand on the idea (e.g., exploring alternatives to ILR for transforming MIDs) and implement CoDA in one of the existing software packages.

## 5. Conclusion

In this work, we introduced compositional isotopic metabolic flux analysis and proposed a method for constructing interpretable ILR transformations of isotopologue distribution simulations and measurements via ordered sequential binary partitions. Our computational experiments showed that compositional I-MFA outperforms the traditional framework, produces more accurate flux estimates and enables computationally inexpensive uncertainty quantification. Ultimately, more accurate fluxotypes directly translate into more reliable predictive models for host-strain optimization and metabolic bottleneck identification.

## Author contributions

A.S.C and T.C. contributed equally to this work as co-authors. A.S.C, T.C. and T.G. carried out the study, developed the mathematical framework, performed the computational experiments, and wrote the original manuscript. N.L.C, T.G. and C.B. contributed to the methodology and visualizations. T.G. and L.K.N. provided supervision and managed the project. All authors contributed to reviewing and editing the manuscript.

## Acknowledgments

This work was funded by the Novo Nordisk Foundation (NNF14OC0009473 and NNF24SA0100980) and by the EU Horizon project DIGIBIO (101060066).

The authors thank Pierre Millard and Sergi Muyo Abad for giving suggestions to this project.

## Data availability

The source code implementing compositional I-MFA, along with all example datasets and reproducible scripts used in this study, are openly available in the GitHub repository at https://github.com/dtuqmcm/cmfa (or via Zenodo DOI: https://doi.org/10.5281/zenodo.21837784).

## Supplementary files

Supplementary file 1: Contains background theory about isotopic metabolic flux analysis and compositional data analysis.

Supplementary file 2: Contains tables showing the metabolic networks and our results.

Supplementary Data 1: Spreadsheet containing results for each of the 10000 measurement simulations from our ground truth flux recovery analysis. This file includes flux estimates and residuals for each run for both traditional I-MFA and compositional I-MFA.

Supplementary Data 2: Spreadsheet containing, for each combination of free fluxes in our sensitivity analysis, the MSE of each flux for traditional I-MFA and compositional I-MFA.

