## Supplementary File 1 for "Improved Metabolic Flux Estimations through Compositional Data Analysis"

### Appendix A. Isotopic metabolic flux analysis

Mathematically, I-MFA considers a metabolic network with $M$ compounds with $I$ total isotopic forms or "isotopomers" and $N$ reactions, with stoichiometric coefficients given by a stoichiometric matrix $\mathbf{S} \in \mathbb{R}^{M \times N}$. Each reaction $k \in \{1, \ldots, N\}$ is assumed to have a known atom transition map, i.e. a bijective mapping $\pi_k : \mathcal{A}_{k,\text{in}} \to \mathcal{A}_{k,\text{out}}$ where $\mathcal{A}_{k,\text{in}}$ and $\mathcal{A}_{k,\text{out}}$ represent the ordered sets of trackable atoms (e.g., carbons) across all substrates and products of reaction $k$. Suppose that each compound $i \in 1, \ldots, M$ has $D(i) \in \mathbb{N}$ isotopologues. Each $j \in 1, \ldots, D(i)$ represents a different mass shift of compound i. The flux vector $\mathbf{v} \in \mathcal{V} \subseteq \mathbb{R}^N$ then determines a labeling pattern $\mathfrak{r}(\mathbf{v}) = (\mathbf{r}_1, \mathbf{r}_2, \ldots, \mathbf{r}_M)$, where each $\mathbf{r}_i \in \mathcal{S}^{D(i)}$ is a point in the $D(i)$-simplex (see section Appendix B for a definition of simplex). For simplicity we assume here that any reaction's flux can in principle be any real number, so that $\mathcal{V} = \mathbb{R}^N$.

The "forward problem" of simulating the label pattern $\mathfrak{r}(\mathbf{v})$ can be solved by writing down a balance equation describing the rate of change of each isotopomer in the network. These equations can be found by combining the atom map and the stoichiometric matrix to produce a matrix $\mathbf{S_I} \in \mathbb{R}^{I \times N}$ of stoichiometric coefficients for isotopomers. At isotopic and metabolic steady state we have $\mathbf{S_I} \cdot \mathbf{v} = 0$, giving $N$ balance equations. Given some known fluxes and isotopomer proportions, other isotopomer proportions can be calculated by solving these equations.

While conceptually simple, this approach to solving the forward problem is computationally demanding due to the large number $I$ of equations that needs to be solved for a realistic model. See Dai and Locasale (2017) for an overview of ways to reduce the number of equations that needs to be solved without loss of information. The most widely-used approach is the elementary metabolite unit (EMU) framework introduced in Antoniewicz et al. (2007).

The "backwards" or "inverse" problem of inferring steady state fluxes from measured isotopomer or isotopologue distributions can be solved by specifying a metric that quantifies discrepancies between labeling patterns, and then solving the forward problem until a flux is found that simulates a label pattern that approximates the measured label pattern sufficiently closely.

The inverse problem can also be formulated as a statistical modeling problem. Following this approach, the key quantity is the probability $p(\mathfrak{r}^{\text{obs}} \mid \mathfrak{r}^{\text{sim}}(\mathbf{v}))$ of observing labeling pattern $\mathfrak{r}^{\text{obs}}$ given a true flux assignment $\mathbf{v}$ with simulated label pattern $\mathfrak{r}^{\text{sim}}(\mathbf{v})$ found by solving the forward problem for $\mathbf{v}$.

#### *Appendix A.1. Sum of squared residuals*

To quantify how closely a simulated label pattern approximates the measured label pattern, most previous implementations of I-MFA have used the weighted sum of squared residuals (SSR). For $n$ measured metabolites, represented as triples of vectors $(\mathbf{r}_i^{\text{obs}}, \mathbf{r}_i^{\text{sim}}, \mathbf{w}_i)$, each with size $D(i)$, the SSR is defined as follows:

$$\text{SSR} = \sum_{i=1}^{n} \sum_{j=1}^{D(i)} \mathbf{w}_{i,j} (\mathbf{r}_{i,j}^{\text{obs}} - \mathbf{r}_{i,j}^{\text{sim}})^2 \tag{A.1}$$

Minimizing SSR is motivated by a connection with the Normal distribution. Assuming unbiased independent and normally-distributed measurement errors, we have

$$\begin{aligned} p(\mathfrak{r}^{\text{obs}} \mid \mathfrak{r}^{\text{sim}}(\mathbf{v}) &= \prod_{i=1}^{n} \mathcal{N}(\mathbf{r}_i^{\text{obs}} \mid \mathbf{r}_i^{\text{sim}}, \sigma_i^2) \\ &= \prod_{i=1}^{n} \prod_{j=1}^{D(i)} \frac{1}{\sqrt{2\pi\sigma_{i,j}^2}} \exp \frac{-(\mathbf{r}_{i,j}^{\text{obs}} - \mathbf{r}_{i,j}^{sim})^2}{2\sigma_{i,j}^2} \end{aligned} \tag{A.2}$$

Taking logs on both sides yields

$$\begin{aligned} \ln p(\mathfrak{r}^{\text{obs}} \mid \mathfrak{r}^{\text{sim}}(\mathbf{v})) &= \sum_{i=1}^{n} \ln \mathcal{N}(\mathbf{r}_i^{\text{obs}} \mid \mathbf{r}_i^{\text{sim}}, \sigma_i^2) \\ &= \sum_{i=1}^{n} \sum_{j=1}^{D(i)} -\frac{1}{2} \ln 2\pi\sigma_{i,j}^2 - \frac{(\mathbf{r}_{i,j}^{\text{obs}} - \mathbf{r}_{i,j}^{sim})^2}{2\sigma_{i,j}^2} \end{aligned} \tag{A.3}$$

Setting $\mathbf{w}_{i,j} = \frac{1}{\sigma_{i,j}^2}$ for each measurement, it is clear that minimizing SSR and maximizing $p(\mathfrak{r}^{\text{obs}} \mid \mathfrak{r}^{\text{sim}}(\mathbf{v}))$ under this measurement model are equivalent.

#### Appendix B. Compositional data analysis

Compositional data consists of units that carry only relative information, so that they can be fully represented by non-negative vectors that sum to a constant $\kappa$ (Pawlowsky-Glahn and Egozcue, 2006). Because the parts of a composition are constrained to a constant sum, they are not mathematically independent; if the values of $D-1$ parts are known, the final component is automatically determined as $\kappa - \kappa \sum_{j=1}^{D-1} x_j$. Consequently, a composition with $D$ parts does not occupy the full $D$-dimensional Euclidean space but instead lies in a restricted $(D-1)$-dimensional sample space known as the simplex. The $D$-simplex $\mathcal{S}^D$ is the set

$$\mathcal{S}^D = \left\{ \mathbf{x} = [x_1, \ldots, x_D] : \forall_{j\in\{1,\ldots,D\}} x_j > 0, \sum_{j=1}^{D} x_j = \kappa \right\} \tag{B.1}$$

In most standard applications, $\kappa = 1$ is used to represent the composition as a vector of proportions.

Figure B.6 illustrates this, showing how a composition in a 3-dimensional Euclidean space maps onto a 2-dimensional simplex $\mathcal{S}^3$.

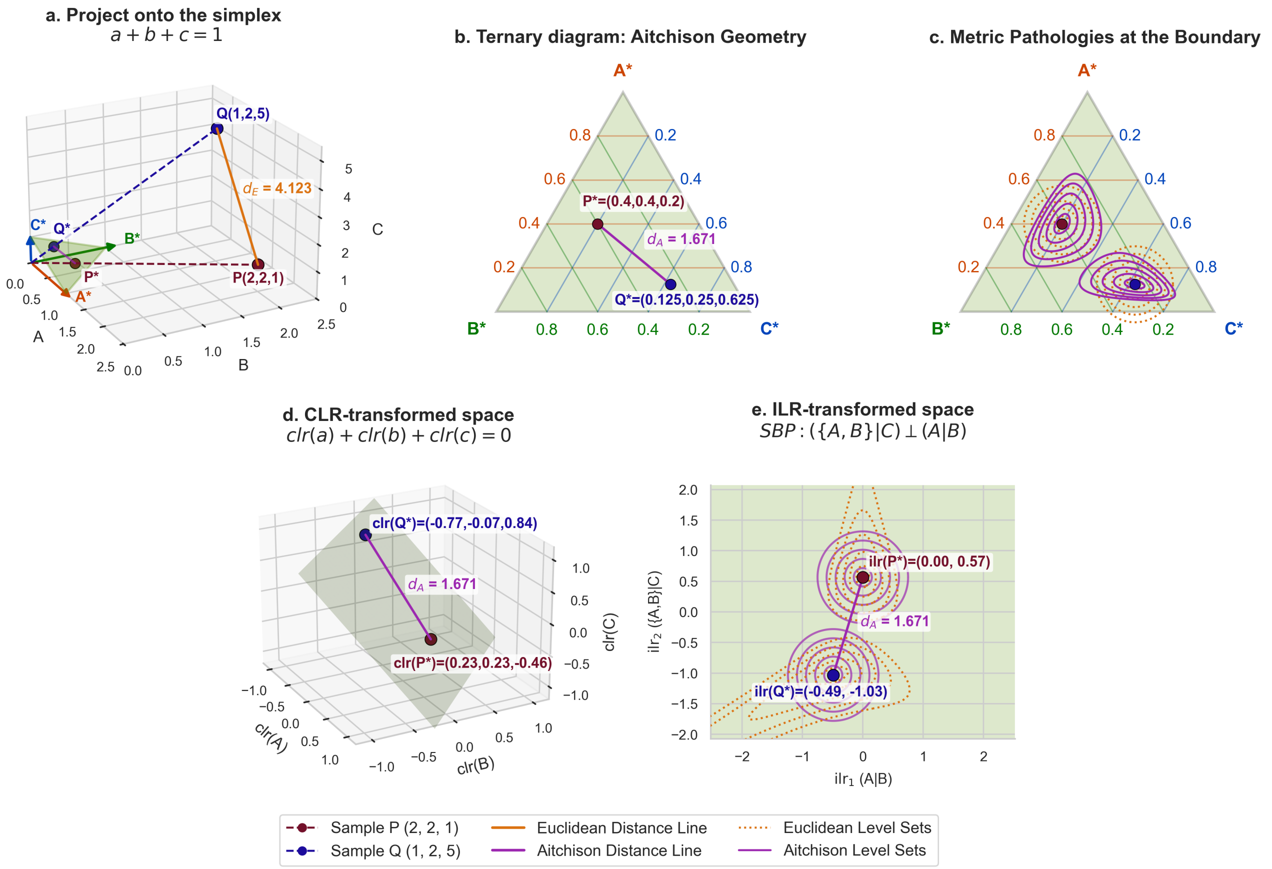


Fig. B.6: **Comparison of Euclidean and compositional (Aitchison) geometries for three-part data** (a) Two 3-component data points $P = (2, 2, 1)$ and $Q = (1, 2, 5)$ and their Euclidean distance ($d_E = 4.123$). The normalization and projection of points P and Q onto the 2-simplex (the constrained ternary plane where $A + B + C = 1$), resulting in the compositional data points P* and Q*. (b) A ternary diagram representing the compositions P* and Q* in Aitchison space. Here, the difference between the points is measured by the Aitchison distance ($d_A = 1.671$), which appropriately scales relative differences rather than absolute ones. (c) An illustration of the metric pathologies that occur when applying Euclidean geometry to compositional data. Orange dotted lines represent level sets of equal Euclidean distance, while purple solid lines denote level sets of equal Aitchison distance. Notably, the Euclidean contours (orange) erroneously exit the simplex, generating mathematically invalid compositions. Furthermore, the Aitchison contours (purple) become highly asymmetrical near the edge of the simplex. (d) The centred log-ratio (CLR) transformation maps each composition to a point in 3-dimensional real space constrained to the hyperplane $\mathrm{clr}(A) + \mathrm{clr}(B) + \mathrm{clr}(C) = 0$ (shaded plane), giving $\mathrm{clr}(P^*) \approx (0.23, 0.23, -0.46)$ and $\mathrm{clr}(Q^*) \approx (-0.77, -0.07, 0.84)$. The Euclidean distance between the two CLR vectors equals the Aitchison distance $d_A$. (e) The isometric log-ratio (ILR) transformation with sequential binary partition(SBP) $\{A, B\}|C \perp A|B$ collapses the 3-part simplex to a 2-dimensional Euclidean space, giving $\mathrm{ilr}(P^*) \approx (0.00, 0.57)$ and $\mathrm{ilr}(Q^*) \approx (-0.49, -1.03)$. Unlike CLR, the ILR representation is unconstrained and bijective, making it directly suitable for standard multivariate statistical methods. The Euclidean distance in ILR space again recovers $d_A$.

It is widely acknowledged that compositional data require specialized analysis methods: see Aitchison (1982) and Pawlowsky-Glahn et al. (2015) for general arguments and Gloor et al. (2017); Quinn et al. (2018); Pawlowsky-Glahn and Egozcue (2006) for discussions of compositional data analysis in specific fields. There are three reasons for this.

First, non-compositional data analysis methods lack principled ways to accommodate the simplex boundaries. For example, naively modeling compositional data with unconstrained statistical models creates a dilemma: either assign probability mass to impossible, non-physical measurements (e.g., negative fractions) or use a sharply truncated distribution. The latter approach leads to misleading uncertainty estimates near the boundaries and causes undesirable behavior (See Figure B.6c) in gradient-based optimizers, which encounter discontinuous loss gradients around the simplex boundaries.

Second, non-compositional methods cannot account for the intrinsic dependencies caused by the sum constraint. If one part of a composition increases, other parts must necessarily decrease, inducing negative correlations. A non-compositional analysis must either ignore these correlations, leading to incorrect results, or attempt to recreate them without explicitly accommodating the underlying compositional structure, which is difficult due to the non-linear relationship between the simplex and Euclidean space.

Third, non-compositional methods typically assume that absolute changes in data are meaningful, whereas compositional data carry only relative information. For example, a shift in labeling fraction from 0.1% to 0.2% represents a 100% relative enrichment (a large physical perturbation), whereas a shift from 50.0% to 50.1% represents a negligible variation. In a standard Euclidean framework, an absolute shift of 0.1 units must be weighted identically regardless of the baseline, distorting the underlying information.

In the following, we review some concepts from compositional data analysis.

*Appendix B.1. Closure*

The closure operation transforms a vector into a composition so that its components sum to a fixed positive real number, $\kappa$. The closure $C(\mathbf{x})$ of a positive vector $\mathbf{x} \in \mathbb{R}_+^D$ with D components is defined as

$$C(\mathbf{x}) = \frac{\kappa \mathbf{x}}{\sum_{j=1}^{D} x_j} \tag{B.2}$$

*Appendix B.2. Log-ratio transformations*

A composition can be mapped from its native simplex to Euclidean space using information-preserving transformations such as the centered log-ratio (CLR) or isometric log-ratio (ILR) transformations (Pawlowsky-Glahn et al., 2015).

Before defining these transformations, it is necessary to introduce the geometric mean, which serves as the central reference component for the composition. For a $D$-part composition $\mathbf{x} = [x_1, x_2, \ldots, x_D]$, the component-wise geometric mean $g(\mathbf{x})$ is defined as the $D$-th root of the product of all its strictly positive parts:

$$\mathrm{g}(\mathbf{x}) = \left(\prod_{j=1}^{D} x_j\right)^{1/D} = \exp\left(\frac{1}{D}\sum_{j=1}^{D} \ln x_j\right) \tag{B.3}$$

By utilizing $g(\mathbf{x})$ as a common denominator, transformations can capture the relative information of the parts while canceling the scaling constant. With this property, the centered log ratio (CLR) transformation maps a $D$ part composition $\mathbf{x}$ with geometric mean $g(\mathbf{x})$ to a real vector of length $D$ according to the following rule:

$$\mathrm{CLR}(\mathbf{x}) = \left[\ln \frac{x_1}{g(\mathbf{x})}, \ldots, \ln \frac{x_D}{g(\mathbf{x})}\right] \tag{B.4}$$

Although the CLR transformation is conceptually intuitive, it yields a $D$-dimensional vector that is subject to a zero-sum constraint $\sum_{i=1}^{D} \mathrm{CLR}(\mathbf{x})_i = 0$. This restricts the transformed data to a $(D-1)$-dimensional subspace, so that CLR-transformed compositions have singular covariance matrices, complicating certain multivariate analyzes.

The isometric log-ratio (ILR) transformation, also known as orthonormal log-ratio (OLR), addresses this issue by lowering the dimension of the output. The key step is to construct an orthonormal basis of the $D$-simplex, that is, a matrix $\mathbf{\Psi} \in \mathbb{R}^{D-1} \times \mathbb{R}^{D}$ whose columns are orthogonal unit vectors. The ILR transformation is then

$$\mathrm{ILR}_{\mathbf{\Psi}}(\mathbf{x}) = \mathbf{\Psi} \cdot \mathrm{CLR}(\mathbf{x}) \tag{B.5}$$

*Appendix B.3. Constructing a basis by sequential binary partition*

Whereas each dimension of $\mathrm{CLR}(\mathbf{x})$ represents the log-ratio of the corresponding composition component to the geometric mean $g(\mathbf{x})$, the dimensions of $\mathrm{ILR}_{\Psi}(\mathbf{x})$ represent log-ratios of the columns of $\mathbf{\Psi}$. The interpretation of $\mathrm{ILR}_{\Psi}(\mathbf{x})$ therefore depends on the choice of orthonormal basis $\mathbf{\Psi}$. A meaningful basis can usually be obtained by constructing a sequential binary partition (SBP). A sequential binary partition of size $D$ is a matrix $\mathrm{SBP} \in \mathbb{R}^{D-1} \times \mathbb{R}^{D}$ whose rows sequentially divide its columns into non-overlapping positive (+) and negative (−) subsets. Each element $\mathrm{SBP}_{ij}$ is either positive with value 1, negative with value −1 or zero, so that $\mathrm{SBP}_{ij} \in \{-1, 0, 1\}$. The first row divides all of the columns; the second row divides either the positive or the negative columns of the first row, with other columns assigned the value zero, and so on. For non-ordered compositional data such as microbial species, there is no principled basis for choosing one particular sequential binary partitioning scheme before knowing which parts are most interesting to compare. For a given sequential binary partition SBP, an orthonormal basis $\mathbf{\Psi}$ can be obtained by normalizing the values of SBP:

$$\mathbf{\Psi}_{ij} = \begin{cases} \frac{1}{r_i}\sqrt{\frac{r_i \cdot s_i}{r_i+s_i}}, & \mathrm{SBP}_{ij} > 0 \\ \frac{1}{s_i}\sqrt{\frac{r_i \cdot s_i}{r_i+s_i}}, & \mathrm{SBP}_{ij} < 0 \\ 0, & SBP_{ij} = 0 \end{cases} \tag{B.6}$$

where $r_i$ and $s_i$ denote the number of positive and negative values in row $i$ of SBP.

*Appendix B.4. Aitchison distance*

The Euclidean distance between two $D$-sized vectors $\mathbf{a}$ and $\mathbf{b}$ is

$$d_E(\mathbf{a}, \mathbf{b}) = \sqrt{\sum_{i=1}^{D}(a_i - b_i)^2} \tag{B.7}$$

Note that, assuming all weights are equal to 1, the Euclidean distance between two vectors is the square root of their SSR.

Aitchison (1982) noted that Euclidean distance does not satisfy scale invariance and sub-compositional coherence, which are desirable properties for a measure of discrepancies between compositions. For an introduction to these desirable properties, see Appendix B.5.

Instead, Aitchison proposed the Aitchison distance $d_A$, defined as follows for $D$-part compositions $a$ and $b$:

$$d_A(\mathbf{a}, \mathbf{b}) = \sqrt{\sum_{i=1}^{D}\left(\ln \frac{a_i}{g(\mathbf{a})} - \ln \frac{b_i}{g(\mathbf{b})}\right)^2} \tag{B.8}$$

Comparing equations B.7 and B.8, it is clear that Aitchison distance is simply the Euclidean distance between the CLR-transformed compositions:

$$d_A(\mathbf{a}, \mathbf{b}) = d_E(\mathrm{CLR}(\mathbf{a}), \mathrm{CLR}(\mathbf{b})) \tag{B.9}$$

Because the ILR transformation defines an isometry between the simplex and Euclidean space, it perfectly preserves geometric distances. Therefore, just as with the CLR transformation, the Aitchison distance between two compositions $\mathbf{a}$ and $\mathbf{b}$ is equivalent to the standard Euclidean distance between their ILR coordinates Pawlowsky-Glahn et al. (2015):

$$d_A(\mathbf{a}, \mathbf{b}) = d_E(\mathrm{ILR}_\Psi(\mathbf{a}), \mathrm{ILR}_\Psi(\mathbf{b})) \tag{B.10}$$

*Appendix B.5. Desirable properties for metrics on compositions*

When working with compositional data analysis principles have to be obeyed. These include scale invariance, subcompositional coherence and permutational invariance. These three principles will be described in the following, and for a deeper explanation, we refer to Pawlowsky-Glahn et al. (2015).

*Appendix B.5.1. Scale invariance*

A function is scale invariant if the variable can be scaled without changing the output of the function, $f(x) = f(\lambda x)$, for every positive real value of $\lambda \in \mathbb{R}^+$. In compositional data, the sum is irrelevant, and only the ratio between the components is important. Therefore, scaling should not influence the analysis result.

*Appendix B.5.2. Permutation invariance*

The order of the components in a composition should not influence the result of the analysis, which is called permutation invariance. An example of a non permutation invariant function is ratio $\frac{A}{B} \neq \frac{B}{A}$.

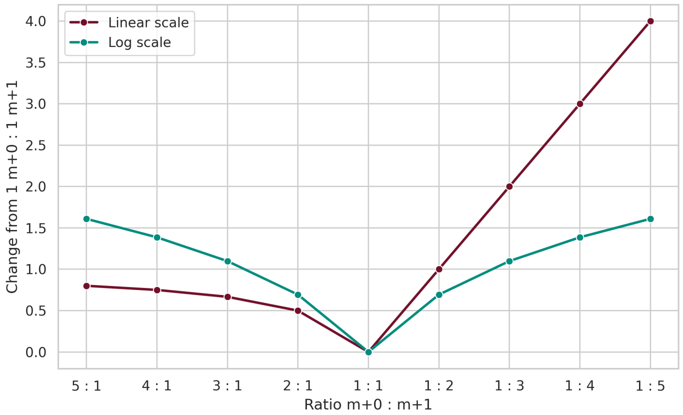


Fig. C.7: **Comparison of Linear and Logarithmic Metrics for Isotopologue Shifts.** The plot illustrates the change in MID measured from a baseline of isotopic equality $(1, m+0 : 1, m+1)$. The linear scale (purple) shows a mathematically inconsistent asymmetry: a shift toward a higher $m+0$ ratio appears significantly larger than an equivalent reciprocal shift toward $m+1$. In contrast, the logarithmic scale (green), grounded in Aitchison geometry, provides a symmetric and scale-invariant measure of change.

*Appendix B.5.3. Subcompositional coherence*

Subcompositional coherence means that the distance between two sub-compositions should be lower than or the same as the distance between the full composition. For a distance metric $d(\cdot, \cdot)$ to be subcompositionally coherent, it must satisfy the following inequality:

$$d(a_s, b_s) \leq d(a, b)$$

where $a_s$ and $b_s$ are subcompositions of $a$ and $b$.

In compositional data analysis, the Aitchison distance strictly satisfies this property. This ensures that the distance between two samples based on a subcomposition is never artificially inflated, guaranteeing that relationships between the retained parts remain logically consistent even when certain components are excluded.

### Appendix C. MIDs are compositions

We first present a simple example that demonstrates how linear metrics fail to preserve symmetric relative changes in I-MFA. Consider the simplest possible MID: a single metabolite with two isotopologues, $m+0$ and $m+1$. This distribution is fully described by the ratio $m + 0 : m + 1$. If a system starts at an isotopic equilibrium of $1 : 1$, a metabolic shift that doubles $m + 1$ yields a $1 : 2$ ratio, while a shift doubling $m + 0$ yields a $2 : 1$ ratio. Chemically, the magnitude of the perturbation from equality is identical in both directions. A linear metric falsely concludes that the $1 : 2$ ratio ($|0.5 - 1.0| = 0.5$) is closer to equality than the $2 : 1$ ratio ($|2.0 - 1.0| = 1.0$). In contrast, as shown in figure C.7, the logarithmic difference $\delta_{\log}(r_1, r_2) = |\ln(r_2) - \ln(r_1)|$ from equality is equal ($\approx 0.69$) for both doubling and halving.

Thus metrics like Aitchison distance that are based on logarithmic differences better match the structure of the underlying problem compared with linear metrics.
