## Supplementary File 2 for "Improved Metabolic Flux Estimations through Compositional Data Analysis"

### Appendix D. Metabolic networks

The reaction stoichiometry, atom transitions and flux allocations are provided in Tables D.1 and D.2 for the toy model and TCA network model.

Table D.1: **Reaction stoichiometry and atom transition of toy network**. [a]For each metabolite, atoms are identified using lower case letters to represent successive atoms of that metabolite. [b]In the columns "Independent flux allocation" the independent fluxes are marked with an △, while a numbers means that the flux-value are fixed in the analysis.

| Reaction number | Reaction stoichiometry | Atom transition [a] | Reaction type | Independent flux allocation |
|---|---|---|---|---|
| 1 | $A \rightarrow B$ | $abc \rightarrow abc$ | Irreversible | 100 |
| 2 | $B \leftrightarrow D$ | $abc \leftrightarrow abc$ | Reversible forward | |
| 3 | $D \leftrightarrow B$ | $abc \leftrightarrow abc$ | Reversible reverse | △ |
| 4 | $B \rightarrow C + E$ | $abc \rightarrow bc + a$ | Irreversible | |
| 5 | $B + C \rightarrow D + E + E$ | $abc + de \rightarrow bcd + a + e$ | Irreversible | △ |
| 6 | $D \rightarrow F$ | $abc \rightarrow abc$ | Irreversible | |

Table D.2: **Reaction stoichiometry and atom transition of TCA cycle**. [a]For each metabolite, atoms are identified using lower case letters to represent successive atoms of that metabolite. An exception is the letter "X", which is used to exclude a metabolite from the isotopomer balance. [b]An △ in Independent flux allocation column is the independent free fluxes while a number denotes the fixed fluxes.

| Reaction number | Reaction stoichiometry | Atom transition [a] | Reaction type | Independent flux allocation [b] |
|---|---|---|---|---|
| 1 | $PYR_{EX} \rightarrow PYR$ | $abc \rightarrow abc$ | Irreversible | 1 |
| 2 | $GLU_{EX} \rightarrow AKG$ | $abcde \rightarrow abcde$ | Irreversible | △ |
| 3 | $PYR \rightarrow ACCOA + CO_2 + NADH$ | $abc \rightarrow bc + a + X$ | Irreversible | |
| 4 | $ACCOA + OAA \rightarrow ICI$ | $ab + def \rightarrow fedbac$ | Irreversible | |
| 5 | $ICI \rightarrow AKG + CO_2 + NADH$ | $abcdef \rightarrow abcde + f + X$ | Irreversible | |
| 6 | $AKG \rightarrow 0.5SUC + 0.5SUC + CO_2 + NADH + ATP$ | $abcde \rightarrow 0.5abcd + 0.5dcba + e + X + X$ | Irreversible | |
| 7 | $SUC \leftrightarrow OAA + FADH_2 + NADH$ | $abcd \leftrightarrow abcd + X + X$ | Reversible Forward | |
| 8 | $OAA + FADH_2 + NADH \leftrightarrow 0.5SUC + 0.5SUC$ | $abcd + X + X \leftrightarrow 0.5abcd + 0.5dcba$ | Reversible Reverse | △ |
| 9 | $OAA \rightarrow PYR + CO_2$ | $abcd \rightarrow abc + d$ | Irreversible | △ |
| 10 | $PYR + CO_2 + ATP \rightarrow OAA$ | $abc + d + X \rightarrow abcd$ | Irreversible | |
| 11 | $2NADH + O_2 \rightarrow 4ATP$ | | Metabolite balance | |
| 12 | $2FADH_2 + O_2 \rightarrow 2ATP$ | | Metabolite balance | |
| 13 | $O_{2\_EX} \rightarrow O_2$ | | Metabolite balance | |
| 14 | $CO_2 \rightarrow CO_{2\_EX}$ | $a \rightarrow a$ | Irreversible | |
| 15 | $ATP \rightarrow ATPM$ | | Metabolite balance | |
| 16 | $PYR \rightarrow PYR_B$ | | Sink | 0.07 |
| 17 | $AKG \rightarrow AKG_B$ | | Sink | 0.23 |
| 18 | $OAA \rightarrow OAA_B$ | | Sink | 0.12 |
| 19 | $PYR + PYR \rightarrow VALX + CO_2$ | $abc + def \rightarrow abefc + d$ | Irreversible | 0.05 |
| 20 | $OAA + PYR \rightarrow LYSX + CO_2$ | $abcd + efg \rightarrow abcdgf + e$ | Irreversible | 0.03 |
| 21 | $OAA \rightarrow ASPX$ | $abcd \rightarrow abcd$ | Isotopomer balance | |

In tables D.3 and D.4 the true reference flux values are shown together with the ranges for the sensitivity analysis of the free fluxes.

Table D.3: True reference flux values and flux range for sensitivity analysis Toy network

| Flux | True value | Value range |
|---|---|---|
| v1 | 100 | |
| v2 | 110 | |
| v3 | 50 | [25, 75] |
| v4 | 20 | |
| v5 | 20 | [10, 30] |
| v6 | 80 | |

Table D.4: True reference flux values and flux range for sensitivity analysis TCA cycle

| Flux | True value | Value range |
|---|---|---|
| v1 | 1 | |
| v2 | 0.201 | [0.1005, 0.3015] |
| v3 | 0.621 | |
| v4 | 0.621 | |
| v5 | 0.621 | |
| v6 | 0.592 | |
| v7 | 0.807 | |
| v8 | 0.215 | [0.1075, 0.3225] |
| v9 | 0.296 | [0.148, 0.444] |
| v10 | 0.475 | |
| v11 | 1.213 | |
| v12 | 0.296 | |
| v13 | 1.509 | |
| v14 | 1.735 | |
| v15 | 5.561 | |
| v16 | 0.07 | |
| v17 | 0.23 | |
| v18 | 0.12 | |
| v19 | 0.05 | |
| v20 | 0.03 | |

Table D.5: True MIDs for Toy network

| Metabolite | m+0 | m+1 | m+2 | m+3 |
|---|---|---|---|---|
| F | 0.0001 | 0.8008 | 0.1983 | 0.0009 |

Table D.6: True MIDs for the TCA cycle. I "-" denotes that the metabolite do not have the corresponding isotopologue

| Metabolite | m+0 | m+1 | m+2 | m+3 | m+4 | m+5 | m+6 |
|---|---|---|---|---|---|---|---|
| lysx | 0.0157 | 0.1171 | 0.2464 | 0.2867 | 0.2135 | 0.1000 | 0.0206 |
| valx | 0.0128 | 0.1025 | 0.2757 | 0.3453 | 0.2115 | 0.0522 | - |
| aspx | 0.0528 | 0.3111 | 0.2970 | 0.2513 | 0.0878 | - | - |
| suc | 0.0753 | 0.4222 | 0.2729 | 0.1769 | 0.0527 | - | - |

### Appendix E. Results as tables

The tables below provide the summary statistics for the estimated fluxes in our ground flux recovery analysis. For each network that we analysed, there is a table for traditional I-MFA and for compositional I-MFA. For each flux the true value is given together with the median, minimum, maximum, 25% quantile, and 75% quantile of the 10000 estimated flux values. Table E.11 compares three confidence interval methods between traditional I-MFA and compositional I-MFA.

Table E.7: **Summary statistics for estimated fluxes in the Toy network for traditional I-MFA.** The table show the independent fluxes and the dependent fluxes. The fixed flux $v1$ is not shown. For each flux, the ground truth value is given. From the 10000 estimated values for each flux the median, minimum, maximum, 25% quantile and the 75% quantile are given.

| Flux | True value | Median | Min | Max | Q25 | Q75 |
|---|---|---|---|---|---|---|
| v2 | 110.000 | 109.845 | 72.563 | 265.517 | 100.731 | 120.876 |
| v3 | 50.000 | 49.986 | 24.443 | 193.630 | 42.848 | 58.922 |
| v4 | 20.000 | 20.015 | 14.012 | 26.861 | 18.889 | 21.168 |
| v5 | 20.000 | 20.015 | 14.012 | 26.861 | 18.889 | 21.168 |
| v6 | 80.000 | 79.985 | 73.139 | 85.988 | 78.832 | 81.111 |

Table E.8: **Summary statistics for estimated fluxes in the Toy network for compositional I-MFA**. The table show the independent fluxes and the dependent fluxes. The fixed flux $v1$ is not shown. For each flux, the ground truth value is given. From the 10000 estimated values for each flux the median, minimum, maximum, 25% quantile and the 75% quantile are given.

| Flux | True value | Median | Min | Max | Q25 | Q75 |
|---|---|---|---|---|---|---|
| v2 | 110.000 | 110.068 | 73.398 | 267.962 | 101.183 | 120.055 |
| v3 | 50.000 | 50.018 | 25.077 | 196.058 | 43.518 | 57.895 |
| v4 | 20.000 | 20.014 | 14.048 | 26.759 | 18.895 | 21.154 |
| v5 | 20.000 | 20.014 | 14.048 | 26.759 | 18.895 | 21.154 |
| v6 | 80.000 | 79.986 | 73.241 | 85.952 | 78.846 | 81.105 |

Table E.9: **Summary statistics for estimated fluxes in the TCA cycle for traditional I-MFA**. The table show the independent fluxes and the dependent fluxes. The fixed fluxes $v1$ and $v16$-$v20$ are not shown. For each flux, the ground truth value is given. From the 10000 estimated values for each flux the median, minimum, maximum, 25% quantile and the 75% quantile are given.

| Flux | True value | Median | Min | Max | Q25 | Q75 |
|---|---|---|---|---|---|---|
| v2 | 0.201 | 0.201 | 0.158 | 0.241 | 0.193 | 0.209 |
| v3 | 0.621 | 0.621 | 0.578 | 0.661 | 0.613 | 0.629 |
| v4 | 0.621 | 0.621 | 0.578 | 0.661 | 0.613 | 0.629 |
| v5 | 0.621 | 0.621 | 0.578 | 0.661 | 0.613 | 0.629 |
| v6 | 0.592 | 0.593 | 0.506 | 0.673 | 0.576 | 0.609 |
| v7 | 0.807 | 0.810 | 0.520 | 2.561 | 0.718 | 0.931 |
| v8 | 0.215 | 0.216 | 0.000 | 1.982 | 0.131 | 0.332 |
| v9 | 0.296 | 0.296 | 0.075 | 0.753 | 0.248 | 0.349 |
| v10 | 0.475 | 0.474 | 0.282 | 0.924 | 0.430 | 0.525 |
| v11 | 1.213 | 1.214 | 1.083 | 1.334 | 1.190 | 1.238 |
| v12 | 0.296 | 0.296 | 0.253 | 0.336 | 0.288 | 0.304 |
| v13 | 1.509 | 1.511 | 1.336 | 1.671 | 1.478 | 1.543 |
| v14 | 1.735 | 1.737 | 1.519 | 1.937 | 1.696 | 1.777 |
| v15 | 5.561 | 5.558 | 4.949 | 6.262 | 5.438 | 5.684 |

Table E.10: **Summary statistics for estimated fluxes in the TCA cycle for compositional I-MFA**. The table shows the independent fluxes and the dependent fluxes. The fixed fluxes $v1$ and $v16$-$v20$ are not shown. For each flux, the ground truth value is given. From the 10000 estimated values for each flux the median, minimum, maximum, 25% quantile and the 75% quantile are given.

| Flux | True value | Median | Min | Max | Q25 | Q75 |
|---|---|---|---|---|---|---|
| v2 | 0.201 | 0.201 | 0.173 | 0.231 | 0.196 | 0.206 |
| v3 | 0.621 | 0.621 | 0.593 | 0.651 | 0.616 | 0.626 |
| v4 | 0.621 | 0.621 | 0.593 | 0.651 | 0.616 | 0.626 |
| v5 | 0.621 | 0.621 | 0.593 | 0.651 | 0.616 | 0.626 |
| v6 | 0.592 | 0.592 | 0.537 | 0.652 | 0.582 | 0.603 |
| v7 | 0.807 | 0.809 | 0.553 | 1.700 | 0.741 | 0.892 |
| v8 | 0.215 | 0.216 | 0.000 | 1.082 | 0.153 | 0.295 |
| v9 | 0.296 | 0.296 | 0.182 | 0.521 | 0.270 | 0.325 |
| v10 | 0.475 | 0.475 | 0.356 | 0.688 | 0.449 | 0.503 |
| v11 | 1.213 | 1.213 | 1.130 | 1.304 | 1.199 | 1.229 |
| v12 | 0.296 | 0.296 | 0.268 | 0.326 | 0.291 | 0.301 |
| v13 | 1.509 | 1.510 | 1.398 | 1.630 | 1.490 | 1.530 |
| v14 | 1.735 | 1.736 | 1.597 | 1.886 | 1.711 | 1.762 |
| v15 | 5.561 | 5.563 | 5.118 | 5.996 | 5.480 | 5.646 |

Table E.11: **95% confidence intervals for all three estimation methods.** The lower bound and upper bound are shown for each free flux in the two example we tested. The asterisk (∗) indicates that the interval reached the predefined bounds of the search space. The bold number is the most narrow CI comparing traditional and compositional I-MFA.

| | Monte Carlo simulation | | Profile likelihood | | Fisher information | |
|---|---|---|---|---|---|---|
| Flux | Traditional | Compositional | Traditional | Compositional | Traditional | Compositional |
| *Toy network* | | | | | | |
| v3 | 3.3E1, 8.5E1 | 3.4E1, 8.0E1 | 9.1E−4∗, 1.1E3∗ | 3.9E1, 1.9E2 | 3.1E−4, 5.9E6 | 2.1E1, 1.4E2 |
| v5 | 1.7E1, 2.3E1 | 1.7E1, 2.3E1 | 5.8E0, 4.1E1 | 1.3E1, 2.8E1 | 3.8E0, 1.1E2 | 1.4E1, 2.9E1 |
| *TCA cycle* | | | | | | |
| v2 | 1.8E−1, 2.2E−1 | 1.9E−1, 2.2E−1 | 8.8E−2, 3.5E−1 | 1.7E−1, 2.4E−1 | 9.7E−2, 4.0E−1 | 1.7E−1, 2.4E−1 |
| v8 | 1.5E−2, 6.9E−1 | 6.3E−2, 5.1E−1 | 9.1E−4∗, 1.1E3∗ | 9.1E−4∗, 1.9E0 | 9.3E−4, 7.2E1 | 3.3E−2, 1.8E0 |
| v9 | 1.7E−1, 4.7E−1 | 2.3E−1, 3.9E−1 | 9.1E−4∗, 5.9E0 | 1.9E−1, 6.1E−1 | 1.1E−2, 5.8E0 | 1.8E−1, 6.2E−1 |
